# Strategy of selection and optimization of single domain antibodies targeting the PHF6 linear peptide within the Tau intrinsically disordered protein

**DOI:** 10.1101/2023.07.18.549252

**Authors:** Justine Mortelecque, Orgeta Zejneli, Séverine Bégard, Nguyen Marine, François-Xavier Cantrelle, Xavier Hanoulle, Jean-Christophe Rain, Morvane Colin, Luc Buée, Isabelle Landrieu, Clément Danis, Elian Dupré

**Affiliations:** CNRS EMR9002 – BSI - Integrative Structural Biology F-59000 Lille, France; Univ. Lille, Inserm, CHU Lille, Institut Pasteur de Lille, U1167 - RID-AGE - Risk Factors and Molecular Determinants of Aging-Related Diseases, F-59000 Lille, France; Univ. Lille, Inserm, CHU-Lille, U1172 - LilNCog - Lille Neuroscience & Cognition, F-59000 Lille, France; Hybrigenic Services, F-91000 Evry-Courcouronnes, France

**Author notes:** Corresponding authors: Dr Isabelle Landrieu, Dr Luc Buée, Dr Elian Dupré. Equal contributions (Orgeta Zejneli, Séverine Bégard).

**Keywords:** single-domain antibody (sdAb, nanobody), antibody engineering, protein-protein interactions, Tau protein (Tau), protein aggregation, structural biology, nuclear magnetic resonance (NMR)

## Abstract

The use of VHHs (Variable domain of the Heavy-chain of the Heavy-chain-only antibodies) as disease-modifying biomolecules in neurodegenerative disorders holds promises including to target aggregation-sensitive proteins. Exploitation of their clinical values dependents however on the capacity to deliver VHHs with optimal physico-chemical properties for their specific context of use. We described previously a VHH with high therapeutic potential in a family of neurodegenerative diseases called tauopathies. The activity of this promising parent VHH named Z70 relies on its binding within the central region of the Tau protein. Accordingly, we carried out random mutagenesis followed by yeast two-hybrid screening to obtain optimized variants. The VHHs selected from this initial screen targeted the same epitope as VHH Z70 as shown using nuclear magnetic resonance spectroscopy and had indeed improved binding affinities according to dissociation constant values obtained by surface plasmon resonance spectroscopy. The improved affinities can be partially rationalized based on three-dimensional structures of three complexes consisting of an optimized VHH and a peptide containing the Tau epitope. Interestingly, the ability of the VHH variants to inhibit Tau aggregation and seeding could not be predicted from their affinity alone. We indeed showed that the *in vitro* and *in cellulo* VHH stabilities are other limiting key factors to their efficacy. Our results demonstrate that only a complete pipeline of experiments, here described, permits a rational selection of optimized VHH variants, resulting in our capacity to propose two VHH variants derived from the parent Z70 for their next development steps.

## Introduction

VHHs (Variable Heavy-chain of the Heavy-chain-only antibodies), also named single domain antibodies or nanobodies (1), are the smallest domains of natural *Camelidae* single chain antibodies that are still capable to recognize an antigen. Their many interesting properties and potential for engineering make VHHs ideal candidates for the design of innovative treatments (2) and new investigation tools (3, 4), including for in-cell imaging (5). Their clinical use has indeed been established in recent years, first with caplacizumab that targets von Willebrand factor to treat thrombotic thrombocytopenic purpura (6) and ozoralizumab that targets TNF-alpha (and human serum albumin) to treat the chronic, autoimmune inflammatory disease rheumatoid arthritis (7). In this later case, the advantages stemming from the biochemical properties of VHHs were put forward, observed as a fast onset of action and the absence of secondary reaction, even in a model of prolonged treatment where no anti-VHH antibodies were formed (8). VHH development covers many additional applications and is gaining ground in neurodegenerative disorders (9, 10) although their medical use in this field has not yet been demonstrated. However, given their better potential than immunoglobulins to be delivered to the central nervous system (11), they are considered as biomolecules of high interest to be developed as disease-modifying next-generation treatments for related pathologies.

VHHs targeting a specific epitope can be generated either through animal immunization (generally in llamas) and screening of the immunized library (12) or directly through the screening of a naïve synthetic library without animal handling (13). VHHs rarely target intrinsically disordered proteins and peptides (14, 15). Nevertheless, VHHs that target the prion protein, alpha-synuclein protein, Aß (amyloid) peptides and Tau protein (16–20), which are all involved in neurodegeneration linked to aggregation, have been reported and shown to block their self-assembly process (21) either *in vitro* or in the cytoplasmic compartment of cells.

VHHs have many interesting biophysico-chemical properties stemming from their low size and unique chain. They present several advantages such as an easy recombinant production in different systems be it prokaryotes, yeasts or insect cells and more (22), good stability and engineering properties (15). This leads to many opportunities to optimize those VHHs for specific use, considering their production (23), their affinity (24), their stability (25) or their capacity of being produced intracellularly (9, 16, 26). In relation to this last point, a significant part of VHHs may be unable to fold properly in the reducing environment of the cytoplasm (4, 27), which inhibits the formation of the disulfide bridges that contribute to the VHH stability, leading to aggregation or degradation (26, 28). Improving intracellular stability and solubility of the VHHs thus requires specific engineering approaches or selection. Indeed, many efforts to engineer intrabody stability or solubility were already invested for VHHs and other types of single domain antibodies through molecular evolution (29), random or targeted amino-acid replacement (30, 31) or as protein fusions (32, 33). These various processes to engineer the VHH physico-chemical properties for specific uses remain highly empirical and pipelines to reach improved properties still need to be better defined.

Given the progress made in the clinical use of VHHs and their interest in neurodegenerative proteinopathies, VHHs have been selected against the neuronal Tau protein to develop new research tools and evaluate their therapeutic potential (16, 20). We initially focused on VHH E4-1 that has the capacity to block *in vitro* Tau aggregation but not Tau aggregation seeding in a cell-reporter model (16). Cell assays show that VHH E4-1 is indeed not stable when expressed in eukaryotic cells. The interest in blocking Tau aggregation in neurons, in a subset of neurodegenerative diseases commonly referred to as tauopathies, including the most prevalent Alzheimer’s disease (AD), led to the optimization of VHH E4-1 for intracellular use, resulting in the selection of VHH Z70. This VHH is able to block Tau aggregation seeding in a reporter cell line and in a ThyTau transgenic murine model of tauopathies (16). Given these promising results showing that VHH Z70 can be used to block Tau aggregation inside neurons and the potential of VHHs to reach nanomolar affinities, further optimization was considered.

We therefore developed an optimization strategy to increase the ability of the lead VHH Z70 to bind its Tau target in the cellular environment. This optimization was challenging given the already high affinity of VHH Z70 for Tau and without compromising binding specificity and other important properties, including conformational stability in cell for intracellular applications. We set-up a pipeline of experiments that all seem important to evaluate the properties of VHHs and succeeded in selecting variant VHHs that retained the optimal properties to ensure their efficiency in blocking Tau seeding in cells while showing an increased affinity for Tau protein.

## Results

### Affinity optimization of VHH Z70

To generate optimized variants of VHH Z70 (hereafter Z70), a strategy of limited random mutagenesis coupled with stringent selection by yeast two-hybrid screening was chosen for binding optimization in intracellular conditions. A cDNA mutant library was thus first built by random mutagenesis, targeting the whole sequence of Z70 to produce a variety of VHH preys (N-terminal GAL4-activation domain fusion, GAL4ADZ70) against the Tau bait (C-terminal LexA fusion, Tau-LexA). The library was transformed in yeast, and screening of the library was carried out by cell-to-cell mating on selective medium. The interaction between the bait (Tau-LexA) and prey (GAL4ADZ70) was detected by the growth of a diploid yeast colony on the selective medium. Growth of this colony, auxotrophic for histidine, is dependent on the transcription of the his3 reporter gene, which requires interaction between the bait and the prey. Mutants of Z70 with an improved affinity for Tau were selected on medium without histidine (His-) by increasing the selection pressure using 3AT (3-amino-1,2,4-triazole), with the latter compound being a competitive inhibitor of the his3 reporter gene product, to reach conditions with limited to undetected interaction of Z70 with Tau (100 mM 3AT). 43 mutants were thus obtained and their sequence analyzed (Fig. S1).

### VHH Z70 mutants sequences selection

Mutants contained 1 to 4 different point mutations resulting in amino acid substitutions and 33 different amino acid positions were found substituted at least once (Fig. 1A). Most substitutions occurred in the framework, with only 1 position in complementarity defining region 1 - CDR1 (T32), 3 in CDR2 (E56, G58, S59) and 1 in CDR3 (P101). Interestingly, 3 positions were highly represented, G115 (23.6% of occurrences, 21 occurrences, always substituted with glutamic acid), R47 (15.7%) and S23 (9%) whereas the others were randomly found between 1 and 4 times (< 5%). Among these 3 positions, S23 was never found mutated alone but mostly in combination with G115. Conversely, G115 and R47 mutations were never found in combination.

**Figure 1.**
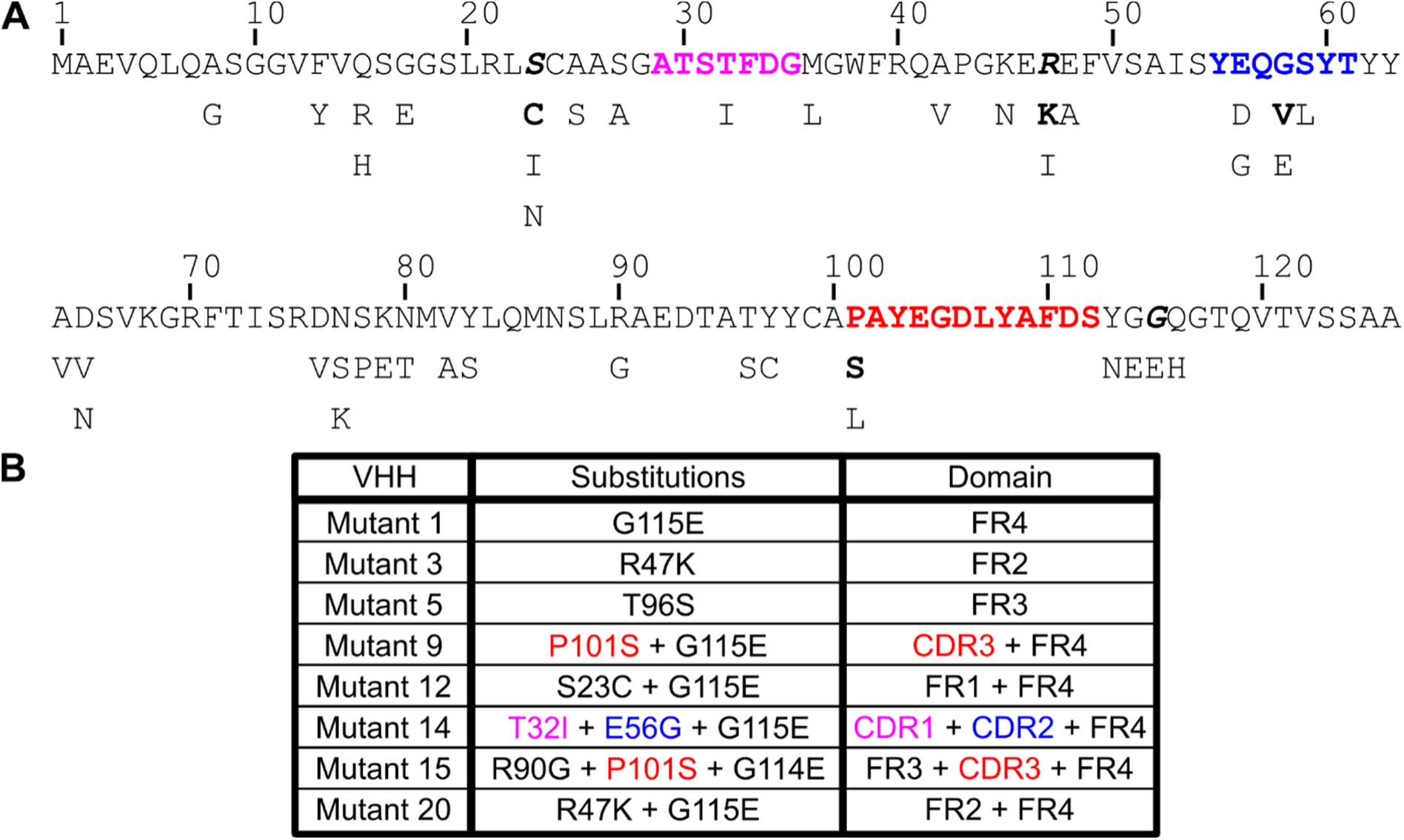
A. Sequence of VHH Z70 and selected substitutions. CDR1, 2 and 3 sequences are represented in pink, blue and red respectively. The 3 amino acids in bold italic are the most frequently substituted. When different substitutions exist that differ in frequency, the most frequent is in bold. B. Overview of the mutants selected for further analysis.

Z70 was already an optimized variant of lead VHH E4-1 as previously described (16), and among the mutations that lead to Z70 only W114G in the FR4 was found further mutated to glutamic acid in this screen (1 occurrence: Mutant 15, G114E in combination with mutations R90G and P101S).

We chose a subset of seven Z70 mutants to represent the variety of mutations, ranging from single to triple mutations and covering framework regions (FRs) and CDRs (Fig. 1B). These selected VHHs had comparable or better interaction with Tau 0N4R than Z70 in yeast two-hybrid system according to one-to-one mating assays (Fig S2). We generated a last mutant, called mutant 20, containing the 2 most prominent mutations G115E and R47K that were not found in combination during the initial screen.

### Affinity measurement

Selection of the optimized mutants was based on their apparent better affinity in a yeast two-hybrid assay compared to Z70. We used surface plasmon resonance (SPR) spectroscopy to confirm and quantify the improved affinities of those mutants using recombinant proteins. Biotinylated Tau 2N4R was immobilized on a streptavidin chip, and the different VHHs were tested as analytes in single cycle kinetics that consist in increasing the analyte concentration in successive passages, without dissociation or regeneration. All eight mutants indeed showed a better affinity than Z70 towards full-length Tau in this assay, further validating the selection process (Fig. 2, S3). Interestingly, the kinetics of the interaction varied depending on the mutant. In most mutants, both k_on_ and k_off_ values were decreased and increased, respectively, except for mutant 5 (T96S) and mutant 15 (R90G + P101S + G114E), which had only either their k_off_ or k_on_ improved respectively. Substitution G115E (mutant 1), which was the most frequent in the initial set of mutants, improved drastically the k_on_ by itself, leading to the best overall affinity (23 nM) measured in this screen. This increased affinity was still observed when combined with additional substitutions in other mutants (9, 12, 14 and 20), but not improved. The k_off_, which might be an interesting parameter as it relates to the time of residence of the VHH on the Tau protein, showed the best improvement for mutants 5 and 20 (R47K + G115E).

**Figure 2.**
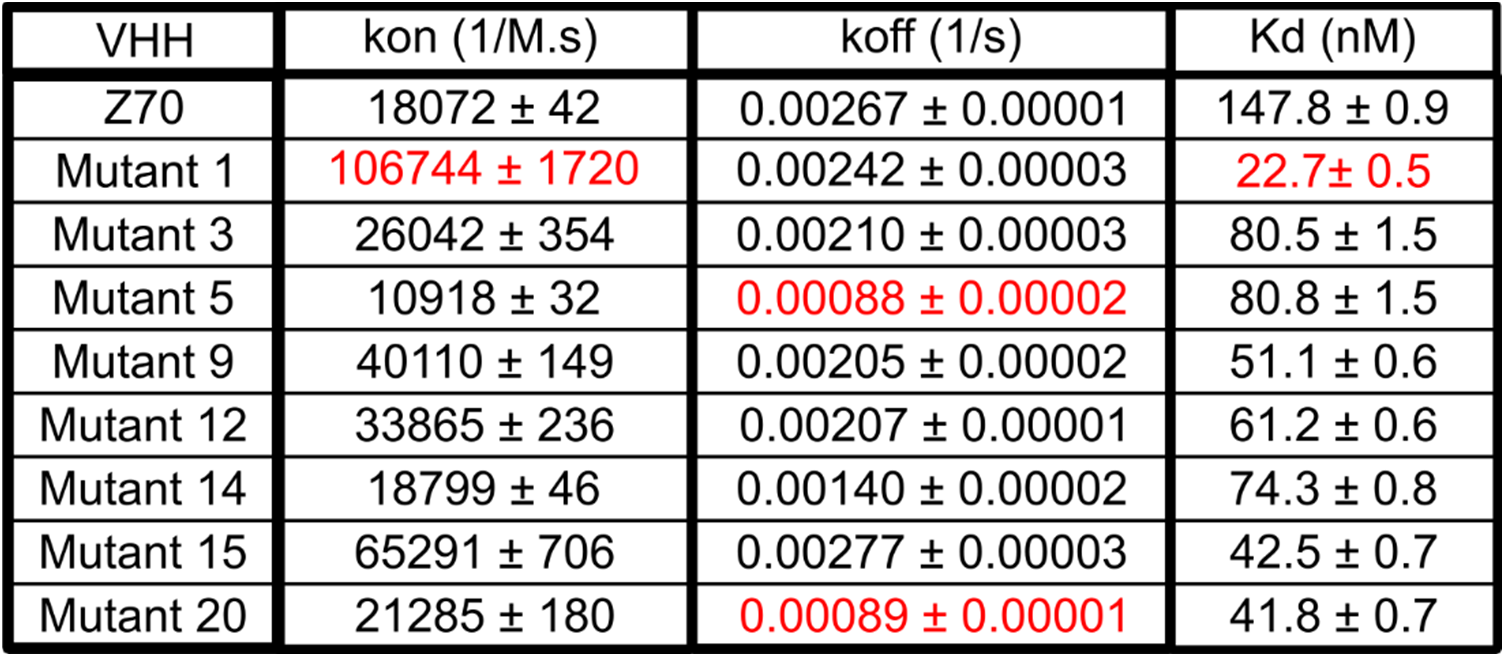
Interaction kinetics between the different VHH and full-length Tau measured by SPR. Best values are highlighted in red.

### Epitope verification

While most amino acid substitutions occurring in the framework regions of the mutants were not expected to modify the Z70 epitope, the optimization screen was carried out to select better binder with full-length Tau bait and modification of the recognition site could thus not be completely ruled out. We first checked whether the VHHs still directly detect Tau using a fluorophore-conjugated VHH. We labeled Z70, mutants 1 (G115E), 3 (R47K) and 20 (double mutation) with TAMRA and used them as probes in Western blots for direct visualization. They were all able to recognize Tau in this setup and their sensitivity did not show drastic changes compared to Z70 (Fig. 3A).

**Figure 3.**
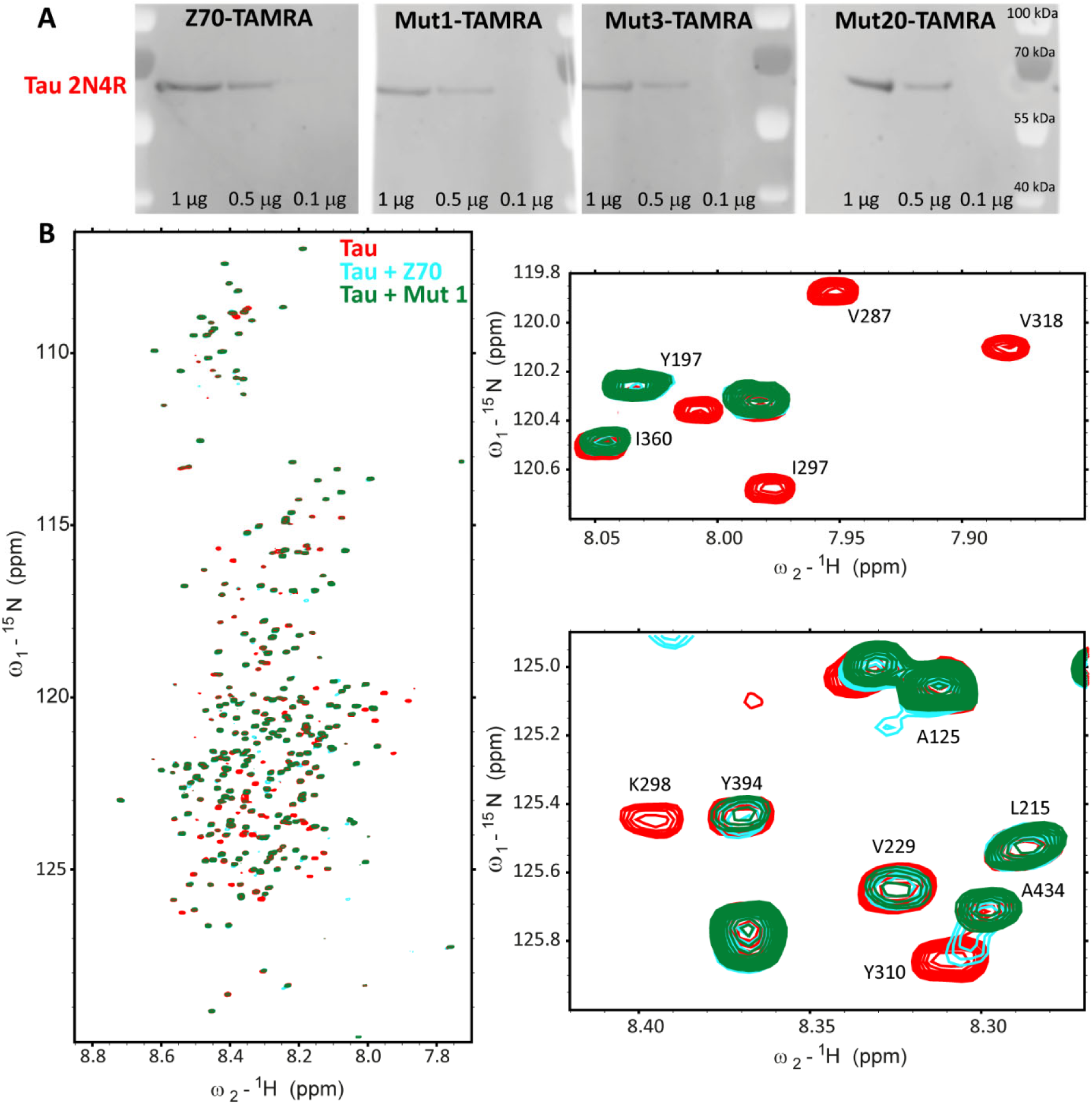
Epitope verification of Z70 mutants. A. Western blot against Tau using TAMRA labeled VHHs. B. Spectrum overlay of Tau alone (in red), Tau with Z70 (in blue, 1:1 molar ratio) or Tau with mutant 1 (in cyan, 1:1 molar ratio).

To obtain further molecular detail on the interaction of the 8 different mutant VHHs, we used ^1^H,^15^N resonance intensity in 2D NMR spectra of Tau as reporters at each amino acid position in Tau sequence (Fig. 3B, S4). The intensity profile is well conserved between the different VHHs with a major loss of intensity for resonances corresponding to residues located in the R3 repeat, similarly to the reported effect of Z70 binding to Tau PHF6 motif (16). As previously reported for Z70, secondary sites of interaction were observed, at the C-terminus of the Tau proline-rich domain and in the R2 repeat, as monitored by the loss of intensity of the corresponding resonances in the spectra of Tau upon interaction (Fig. S4). Superposition of the Tau spectra obtained for each VHH revealed subtle differences that might be due to differences in the kinetics of the interaction between Tau and the VHHs because resonance intensity is not strictly related to binding but also to local dynamics and kinetics of the interaction (34) (Fig. 2). Even for mutant 9, which has the P101S mutation within the CDR3 recognition loop, the perturbations of the resonances were similar to those observed upon Z70 binding to Tau, confirming the conservation of the PHF6 motif inside R3 as the main interaction site of the whole series of VHHs. For mutant 15 (triple mutation), which also contains the P101S mutation within the CDR3, addition to Tau led to extended loss of intensity for many Tau resonances. This was attributed to the formation of aggregates in the conditions of this assay that prevented definitive confirmation of the conservation of the Z70 epitope.

### Effect of substitutions on the structure of mutant VHHs in complex with their peptide epitope

To rationalize the potential effect of the mutations on the interaction, and particularly the effect of the most abundant mutations in the screen, G115E and R47K, we have solved the structures of mutants 1 (G115E), 3 (R47K) and 20 (G115E + R47K) in complex with a Tau[301-312] peptide by X-ray crystallography. We previously reported the crystallographic structure of Z70 in complex with the same peptide containing the PHF6 motif (16) (pdb code : 7QCQ), allowing a straightforward comparison. The structures obtained at a resolution of 1.83 Å, 2.35 Å and 1.54 Å for mutant 1, 3 and 20, respectively, showed no major conformational differences with the parent Z70 bound to its epitope with backbone RMSD values of 0.517 Å between mutant 1 and Z70, 0.545 Å between mutant 3 and Z70 and 0.455 Å between mutant 20 and Z70. (Fig. 4). The interaction between these VHHs and the PHF6 Tau peptide were conserved and involved the formation of a short intermolecular β-sheet between the VHH CDR3 and the PHF6 peptide as previously described for Z70 in (16) (Fig. 5A). G115 in Z70 is in a flexible loop (from F110 to G117) that is partially not resolved in the structure (from D111 to G115) whereas E115 electron density in mutant 20 was well defined. Conversely, 3 residues of this loop (S112-Y113-G114) remained unresolved in the absence of an electron density. Similarly, in mutant 1, this loop remained partially unresolved but some partial electron density can be observed for E115, even if not sufficient to permit a residue replacement. Mutant 3 did not show any more electron density than Z70 for G115, illustrating the importance of the E115 substitution in tightening this loop (Fig. 5B). Positioning of the side chains of residues E115 and K47 in close proximity in mutant 20 suggested the formation of a salt bridge that could tether part of the CDR3 long loop to the core of the VHH.

**Figure 4.**
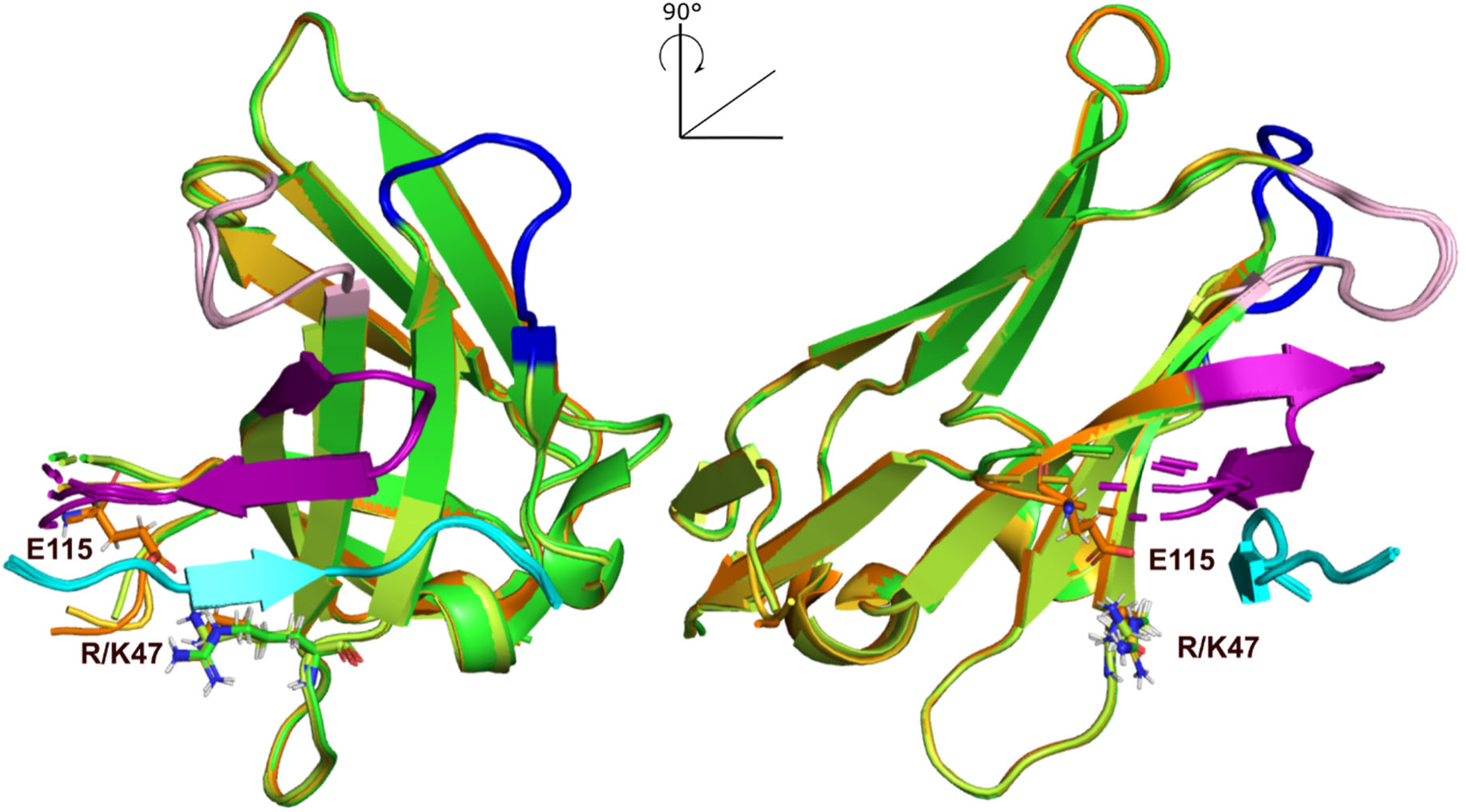
Comparison of Z70, mutant 1, 3 and 20 structures. Superposition of Z70 (in green, pdb 7qcq), mutant 1 (in gold, pdb 8opi), mutant 3 (in pale green, pdb 8pii) and mutant 20 (in orange, pdb 8op0). Their CDR1 are in pink, CDR2 in blue and CDR3 in purple. Tau PHF6 peptide is colored cyan.

**Figure 5.**
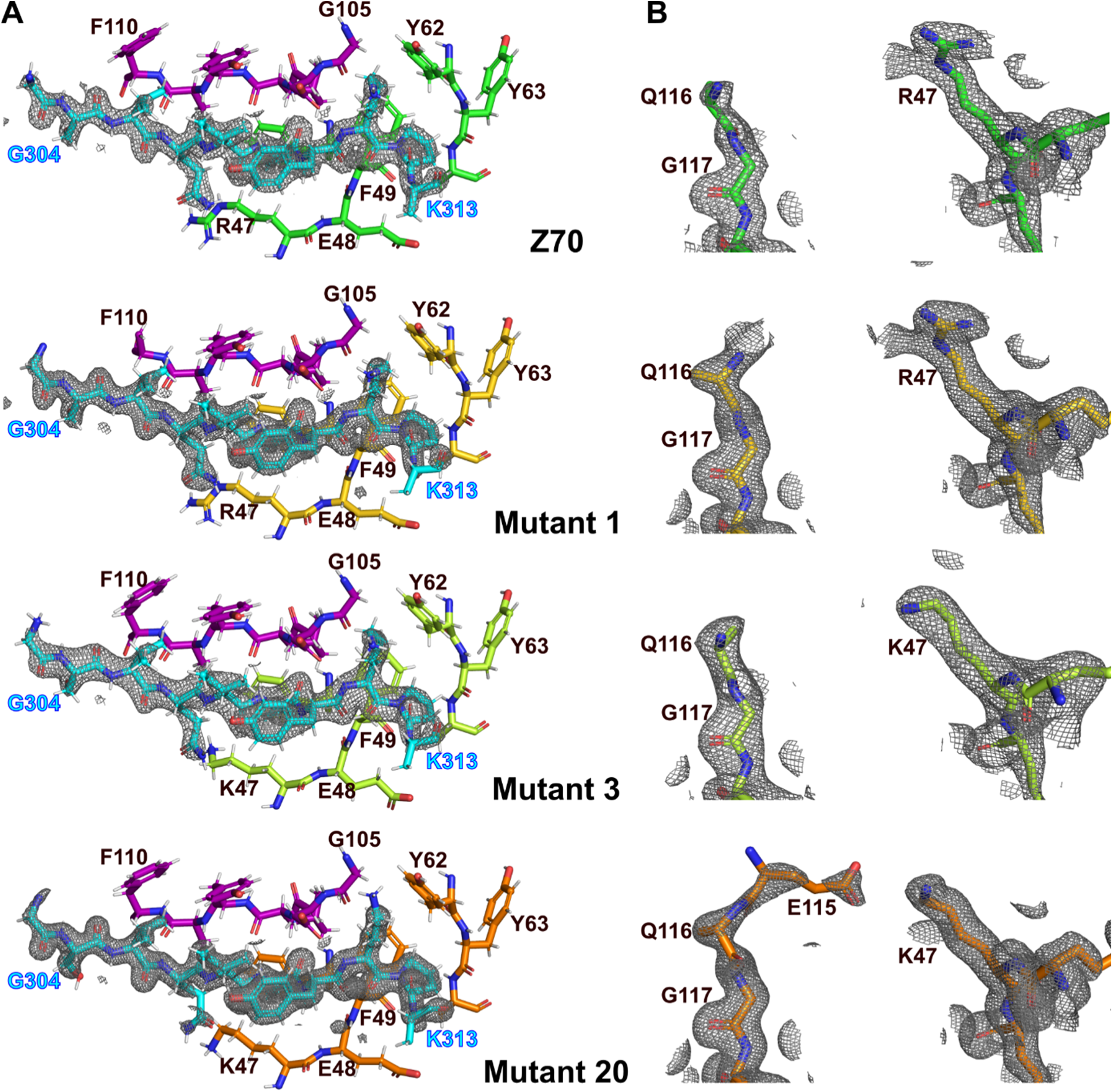
Comparison of Z70, mutant 1, 3 and 20 electron density maps. Z70, mutant 1, 3 and 20 are represented in green, gold, pale green and orange, respectively. Tau PHF6 peptide is colored cyan. A. Environment of the peptide in all 3 structures remains unchanged. B. Electron density (2fo-fc map contoured at 1 σ), shown as mesh representation, corresponding to the region surrounding residues 115-117 (left) and 46-48 (right) in VHH Z70 (top), mutant 1 (middle) and mutant 20 (bottom), respectively, showing the appearance of density for residue 115 from Z70 to mutant 20.

### VHH thermal stability

To verify that the affinity optimization of the Z70-derived VHHs had not compromised their stability, we measured their *in vitro* melting temperatures (Tm). The Tm was measured using a fluorescent reporter whose fluorescence increases once it binds the hydrophobic regions exposed during denaturation. Z70 had a Tm of 59.9 ±0.9°C that falls into the range reported for single domain antibodies (25). Although there were few sequence differences between the individual VHH mutants, their thermal stability temperature range spanned about 15°C. The G115E substitution in mutant 1 had little effect on VHH stability, with a loss of thermal stability of around 2°C (Fig. 6). On the contrary, the P101S and S23C substitutions destabilized the VHHs, as shown by the low melting temperatures of mutants 9, 15 (both carrying P101S, with denaturation temperature dropping by 10°C) and of mutant 12 (carrying S23C, with denaturation temperature dropping by 7°C). Although these mutations provided an increased affinity compared to Z70, the observed destabilization might contribute to a decrease efficiency in functional assays.

**Figure 6.**
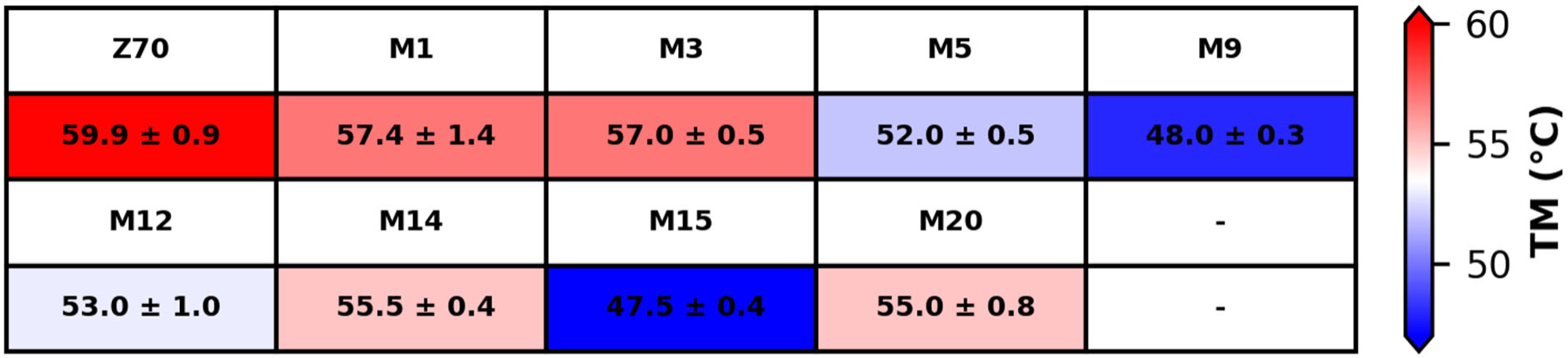
Thermal stabilities of the different VHH in NMR buffer.

### Self-aggregation of the VHH-mCherry constructs in HEK293 cells

We expect the VHH series derived from Z70 to present properties allowing their use in intracellular applications, similarly to the parent VHH. VHHs have already been shown to self-aggregate in cellular environments (26) and we showed that the purified mutated VHHs present a range of *in vitro* thermal stabilities that suggest their self-aggregation tendency in the intracellular compartment might also be affected (Fig 6). We thus used HEK293 cells transfected with pmCherry constructs that produced VHH-mCherry to evaluate the aggregation of the Z70-derived VHH series expressed in the cellular environment. The presence of aggregates is noticed by the appearance of “puncta” inside the cells in contrast with a uniform fluorescence across the cell in the absence of aggregates as already described (26) (Fig. S5). Frequencies of self-aggregation of the various VHH-mCherry fusions were variable. The parent Z70 as well as mutant 1 (G115E), 3 (R47K) and 12 (S23C + G1115E) showed low aggregation propensity, with puncta appearing in less than 10% of the transfected cells (Fig. 7A). Other mutants showed higher aggregation propensity, such as mutants 5 (T96S), 9 (P101S + G115E) and 15 (R90G + P101S + G114E) with puncta in more than about half of the cells. This intracellular aggregation propensity is well correlated to the melting temperature (Pearson correlation coefficient of −0.87; Fig. 7B). Nevertheless mutants 5 and 12 that had similar melting temperatures (51.5-54°C) showed different in-cell behaviors with high aggregation propensity for mutant 5 (about 50% of cells with puncta) and low for mutant 12 (about 10% of cells with puncta).

**Figure 7.**
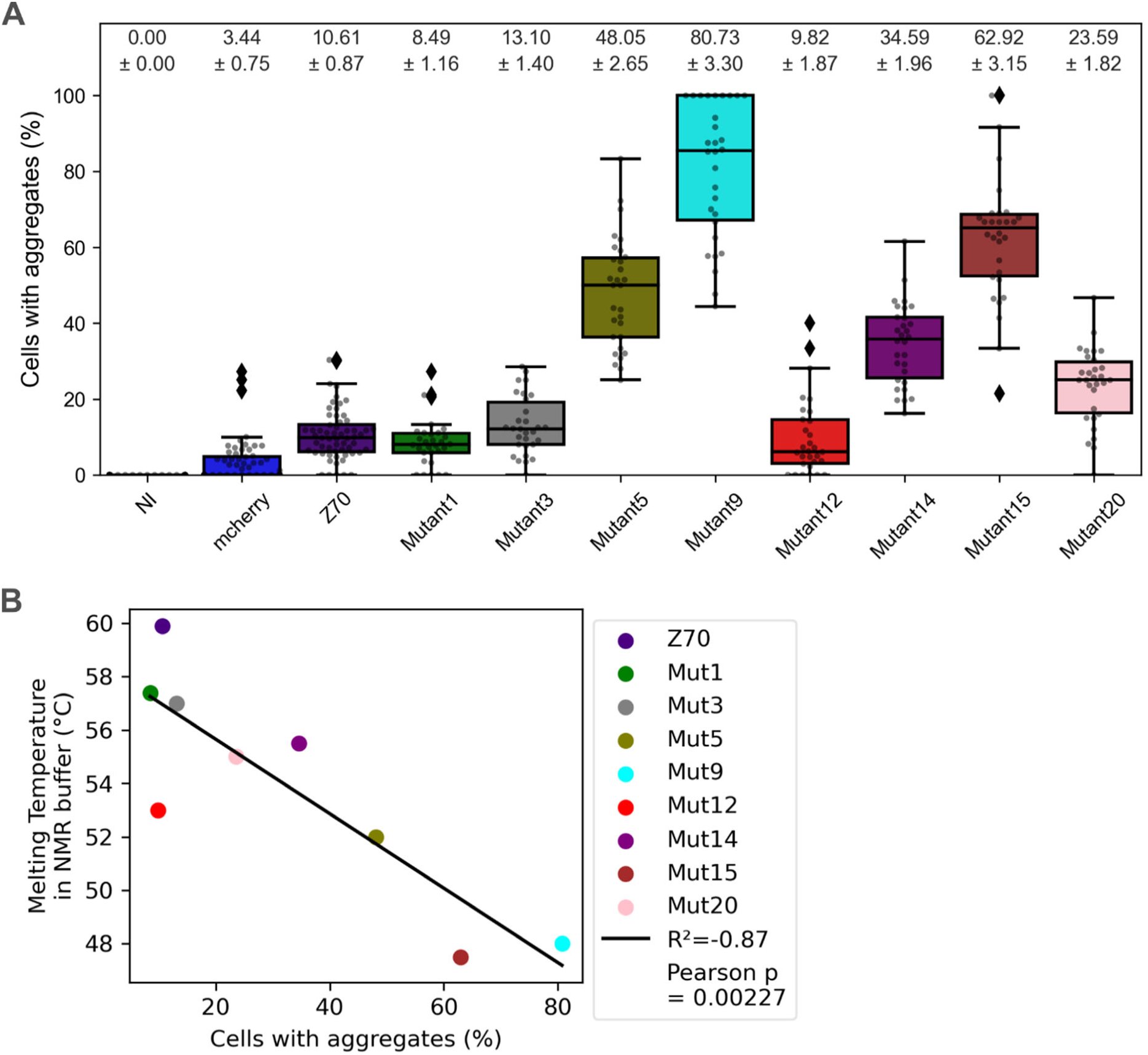
In-cell stabilities of the VHHs. A. Percentage of mCherry positive HEK293 cells showing aggregates (puncta) of VHH-mCherry 48 hours after transfection. B. Pearson plot of the percentage of cells with aggregates in the self-aggregation assay versus melting temperatures in NMR buffer.

### Inhibition of *in vitro* Tau aggregation

Z70 was primarily selected for its ability to inhibit Tau aggregation both *in vitro* and *in vivo* (16). Z70 shows a very potent inhibition of *in vitro* Tau aggregation in the heparin mediated assay in the presence of thioflavin as a fluorescent reporter of the aggregation. We therefore used this assay to evaluate the efficiency of the different mutants to inhibit Tau aggregation in comparison with Z70 at sub-equimolar ratios (Fig. S6, S7). To facilitate comparison between the different VHHs, the data from the different aggregation tests were analyzed as a ratio of Thioflavin T fluorescence intensity in the presence of VHH compared to the positive control (Tau alone) for all points in the dynamic range of fluorescence corresponding to an intensity between 10 and 90% of the maximum fluorescence obtained for the positive control, in the absence of any VHH.

When using a VHH:Tau ratio of 0.5:1 (Fig. S6), we could observe that mutants 1, 3, 12, 14 and 20 seemed to be more efficient than Z70, reducing the observed fluorescence up to 91% for mutant 3 compared to 82% for Z70. In contrast, mutants 5 and 15 were less effective (75 or 68% reduction) and mutant 9 was performing poorly (Fig. 8A), with only 27% reduction, which was mainly due to its tendency to self-aggregate in the condition of the present test, giving rise to higher fluorescence intensity than Tau alone (Fig. S6, S7). The use of a VHH:Tau ratio of 0.2:1 (Fig. S7) showed some differences in these tendencies but confirmed mutants 3 and 20 to be the most efficient, with 75% reduction (Fig. 8B). Surprisingly, mutant 5 repeatedly showed a very good effect (75%) at this low ratio, revealing the understatement of its efficacy due to its propensity to self-aggregate, as seen for mutant 9 (Fig. S7).

**Figure 8.**
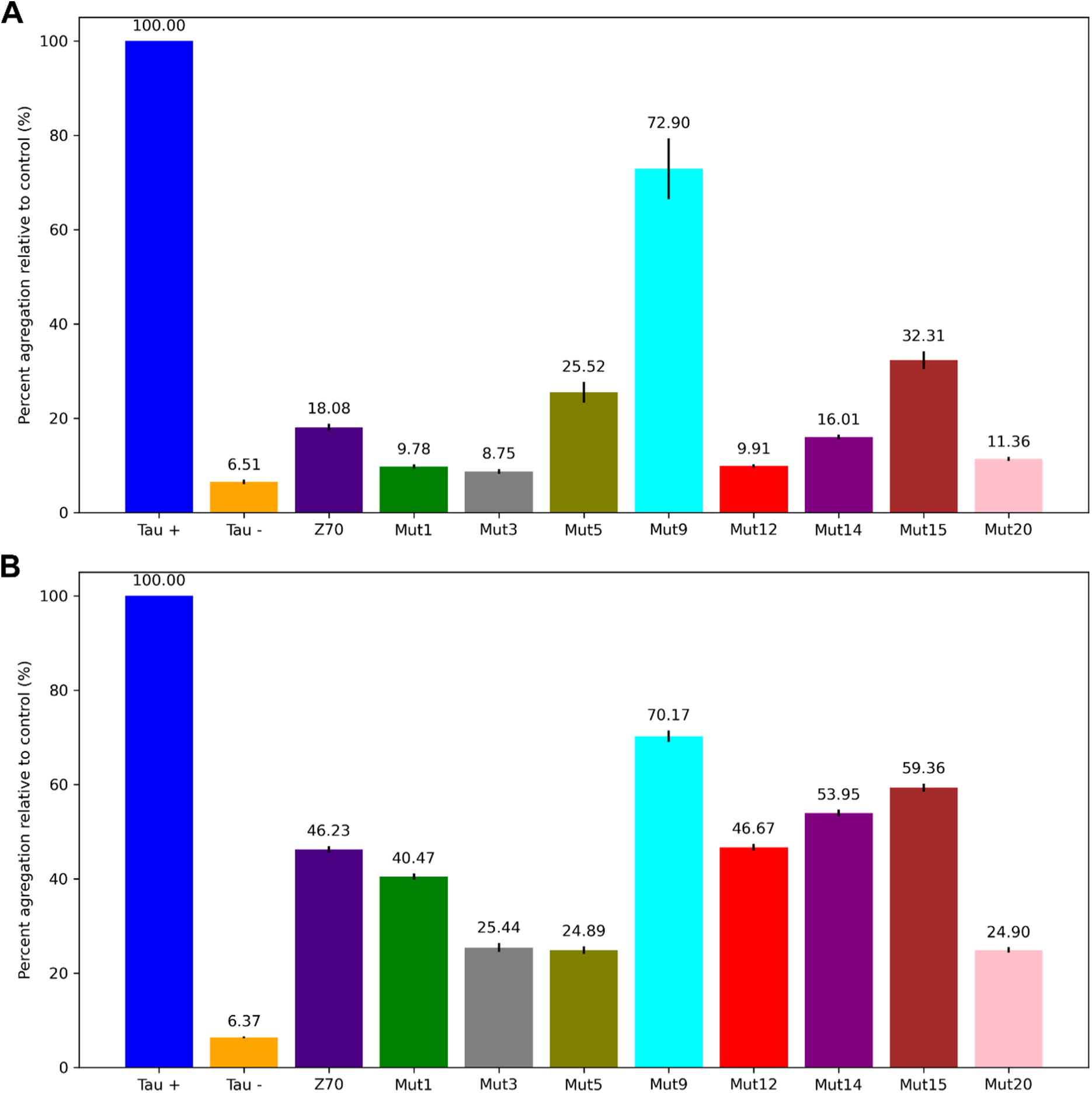
*In vitro* inhibition of Tau aggregation by the VHHs. Positive and negative controls correspond to Tau in the presence of heparin (Tau +) and the absence of heparin (Tau -), respectively. The other conditions correspond to Tau in the presence of heparin and the different VHHs at a VHH:Tau ratio of A. 0.5:1 or B. 0.2:1. The percentage of aggregation corresponds to the fluorescence intensity ratio between the different conditions and Tau positive control (Tau +) during the aggregation process. Data are represented as mean ± sem, the mean value is written above the bars.

### Inhibition of Tau seeding in HEK293 Tau repeat domain (RD) P301S FRET Biosensor reporter cells

We previously described the efficient inhibition of Tau seeding by Z70 in a cellular model (16). As the various mutants had different inhibitory effects in the *in vitro* Tau aggregation assay, we checked their efficiency in this cellular model as a comparison to and validation of the *in vitro* assay. The HEK293 seeding reporter cell line model constitutively expresses Tau RD (corresponding to Tau MicroTubule Binding Domain, MTBD or Tau[244-372]), with a P301S mutation, fused to either CFP (Cyan Fluorescent Protein) or YFP (Yellow Fluorescent Protein) that together generate a FRET (Forster Resonance Energy Transfer) signal upon MTBD-P301S aggregation seeding (35). The intracellular aggregation of MTBD-P301S protein is induced by treating the cells with Tau seeds (heparin-induced MTBD fibrils). VHHs were transfected one day prior to MTBD seed treatment. VHH F8-2 is used as negative control since its binding site is outside the MTBD as described previously (16, 20).

VHH F8-2 negative control was used as a 100% FRET positive reference in the mCherry gated population for each individual experiment to minimize discrepancies arising from transfection efficiencies. Mutants 1, 3 and 12 retained at least Z70 level of efficiency (60.6 ± 1.7 % of the reference) in this assay (Fig. 9) with 63.9 ± 1.9 %, 62.8 ± 2.8 % and 59.4 ± 1.3 % respectively, with mutant 20 following closely (68.8 ± 3.7 %). However, none of these mutants revealed significant difference when compared to Z70. Mutants 5 and 14 gave lower inhibition of seeding (82.3 ± 5.2 % and 81.3 ± 3.4 %) while mutants 9 and 15 failed to inhibit the seeding.

**Figure 9.**
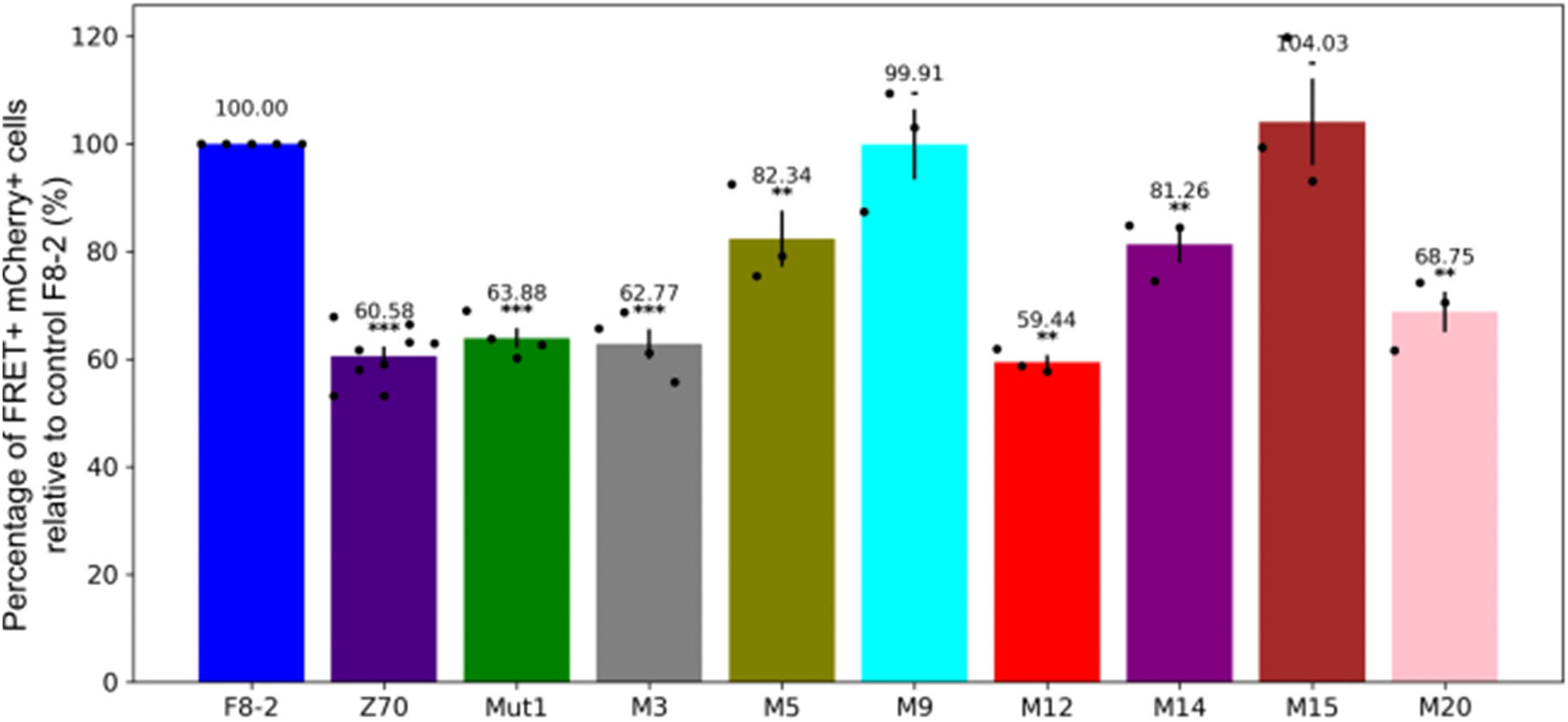
*In cellulo* inhibition of Tau seeding by the VHHs. The percentage of FRET positive cells, corresponding to cells with aggregated Tau, is given for the mCherry-gated population, corresponding to transfected cells. Values are normalized relative to the experiment with F8-2 VHH-mCherry. Data are represented as mean ± sem, the mean value is given above the bars and individual points are plotted. Statistical comparison between each VHH and F8-2 was carried out, *** is p < 0.001, ** p < 0.01 and – is not significant.

### Selection of Z70 VHH variants based on complementary assays

We finally used the complementary *in vitro* and cellular assays to select Z70 VHH derived mutants with a global view on their properties.

Interestingly, there is a correlation between the *in vitro* aggregation assay and the VHH inhibition capacity in the cellular seeding assay (Pearson correlation coefficient of 0.78; Fig. 10A) with mutant 9 being a clear outlier (Pearson correlation coefficient of 0.87 without mutant 9; Fig. 10A). The VHHs with lower performance in the *in vitro* assay performed poorly in the cellular assay (e.g. mutant 15). The mutant 5, 9 and 15 poor *in vitro* inhibition efficiencies were revealed by low thermal stabilities, with low denaturation temperatures between 47.1°C and 52.5°C, coupled to poor stability in cells, with 45.4 to 84% of cells transfected with mCherry-VHH showing puncta. However, some VHHs retaining an inhibition efficiency close to parent Z70 in *in vitro* aggregation assays did not necessarily perform well in the cellular assays (e.g. mutant 14). It is indeed striking that aggregation propensity of the VHH inside the cell is highly correlated to the inefficiency of the VHH to inhibit seeding in the reporter cells (Pearson correlation coefficient of 0.96; Fig. 10B). At the other end of the inhibition efficiency spectrum, parent Z70 as well as mutants 1, 3 and 12 showed low aggregation propensity in eukaryotic cells, with puncta appearing in less than 10% of the transfected cells (Fig. 7A) and were efficient inhibitors in the cellular seeding assays, the latter being consistent with their availability in the cytoplasm. Of note, mutant 12 despite its efficiency in these assays had a relatively low thermal stability (denaturation temperature 53°C, Fig. 7B).

**Figure 10.**
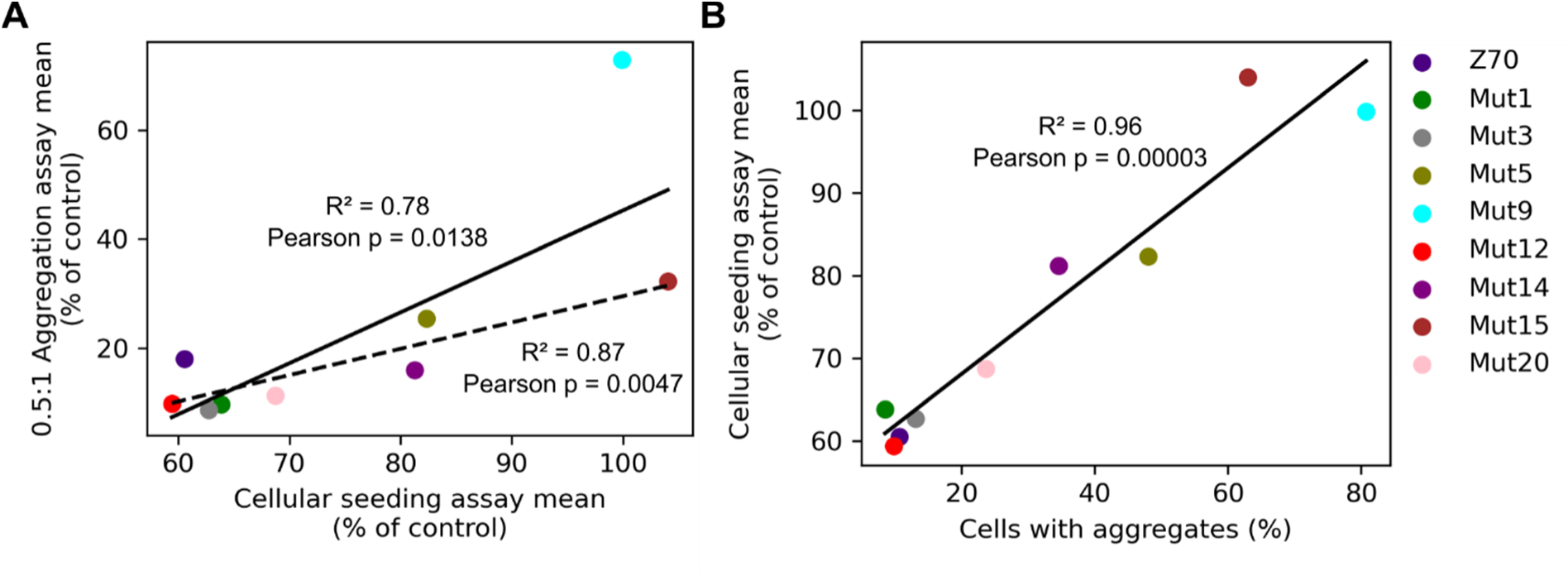
Pearson correlation plots of: A. mean values of percentage of Tau seeding in HEK293 Tau repeat domain (RD) P301S FRET Biosensor seeding reporter cells (Cellular seeding assay) versus mean values of *in vitro* aggregation assay with a ratio of 0.5 VHH for 1 Tau. Dashed line is omitting the outlier Mutant 9. B. Mean values of percentage of Tau seeding in HEK293 Tau repeat domain (RD) P301S FRET Biosensor seeding reporter cells (Cellular seeding assay) versus percentage of cells showing punctate aggregates in HEK293 cells

## Discussion

Optimization of Z70 that binds a linear sequence within the intrinsically disordered Tau protein led to the selection of 8 mutants. All mutants were indeed optimized in their affinity towards the initial epitope, up to more than 6-fold. However, some mutants lost some other essential properties in term of stability when produced in bacteria periplasm and purified and/or when expressed in eukaryotic cells. Yet, they were selected in the two-hybrid screen as better binders. Fusion to the GAL4-activation domain might help to stabilize these VHHs. However, our additional screen of in-cell stability is also done with a fusion with the mCherry protein, which did not rescue the poor stability of some of the mutants, showing that this is an unlikely explanation. In the yeast 2 hybrid assay in these screening conditions, the expression and the stability do not seem to be limiting points. This might not be the case in the seeding reporter cells, in which Tau RD is expressed at high level and blocking the seeding thus requests optimal stability and binding properties of the VHHs.

The affinities that we have reached with the Z70 VHH series might still seem moderate considering the ability of some VHHs to bind to their target with sub nanomolar affinities (36, 37). However, VHHs are particularly efficient to bind cavities as epitopes (38), which is not the case when binding a linear Tau epitope. Moreover, only a few VHHs directed against peptides are described, and in particular one was developed as a capture and detection tool thanks to its high affinity defined by a Kd of 1.4 nM (39). Interestingly, similarly to the interaction of Z70 and the PHF6 peptide of Tau, this peptide engages in the formation of an intramolecular β-sheet with the VHH recognition loop, which might underly the stability of the complex.

The two most common affinity enhancing substitutions (G115E and R47K) were never selected together in this initial screening leading us to generate such a mutant. Even though mutants 1 and 3 respective *in vitro* affinities were confirmed to be better than Z70, mutant 20 failed to give better affinity parameters as a combination of the 2 substitutions, suggesting that the mechanisms by which these substitutions enhance the affinity are not complementary. Atomic resolution structures of mutant 1, 3 and 20 combined with kinetics parameters of the interactions also suggest that their mechanisms of affinity enhancement might indeed differ. Mutant 1 with the introduction of a glutamic acid residue at position 115 had an improved k_on_ and very little structural difference with Z70, which argue for a charge effect facilitating the interaction with the basic Tau protein (40, 41). The same effect might be achieved through the G114E substitution in mutant 15 that also had an increased k_on_ compared to Z70. Mutant 20 structural data suggest that the mutated residues G115E and R47K could form a salt bridge that would partially restrict the CDR3 flexibility as E115 is located at the loop basis, and potentially decrease the entropic cost of the interaction. The ensuing stabilization of the complex is supported by the lowest k_off_ of the VHH series for mutant 20. The link between a decrease flexibility of the CDR3 loop and a stronger antigen interaction is also demonstrated by the presence of an extra-disulfide bond between the CDR3 and the CDR1 present in many VHHs (42). The additional charges of mutant 1 might account for its excellent intracellular solubility (28) whereas the double mutation by neutralizing the positive charge at position 47 lead to a decrease net charge that could result in the poorer solubility in cells of mutant 20.

The correlation between the thermal stability of the VHHs and their in-cell stability was very good (Fig. 7B), with the noticeable exception of mutant 12 (S23C + G115E) that has a lower thermal stability than expected from all the other assays. We have found a very good correlation between the *in vitro* aggregation assay and the cellular seeding assay (Fig. 10), at the exception of mutant 9 (P101S + G115E). This can be easily rationalized based on the poor in-cell solubility and thermal stability of mutant 9. In a general manner, VHHs with poor intracellular solubility (mutant 5, 9, 14, 15, Figs. 7A and 10B) were found incapable of preventing intracellular Tau seeding while more soluble VHHs had better results (Figs. 9 and 10B). Our experiments thus agree with previous research suggesting that for single chain antibodies scFv the intracellular efficiency is dictated by its intracellular stability rather than its *in vitro* affinity for its target (28).

Finally, although some optimized mutants were able to better reduce aggregation in the *in vitro* assay than Z70 and show good intracellular activity, we did not replicate this improvement in the seeding reporter cells. This led us to hypothesize that for these mutants, we have reached a limit of inhibition in the seeding reporter cells and are unable to differentiate however gains were to be had with the mutants, and further *in vivo* validation would be of interest.

We found that the *in vitro* experiments were reasonably predictive of the cellular assay outcomes and could serve as a first step of selection. Outliers in the correlations however argue that only the full set of complementary experiments can offer a comprehensive view of the VHHs properties. Among the different assays, the VHH self-aggregation assay in HEK293 cells provided the best correlation with other tests and particularly the cellular Tau aggregation assay, making it a very good VHH assessment tool when targeting intracellular activity. Among the different optimized mutants, mutants 1 (G115E) and 3 (R47K) showed the best behaviors in the different assays conducted here and could be good candidates for further testing in *in vivo* experiments as improved versions of Z70.

### Experimental Procedures

#### Optimization of VHH Z70 affinity

VHH Z70 was amplified from pHEN2 plasmid using Taq polymerase with 14 mM MgCl_2_ and 0.2 mM MnCl_2_ and a modified nucleotide pool (43). The amplified cDNAs were transformed in yeast Y187 strain, together with a digested empty derivative of pGADGH vector (44), allowing recombination by gap repair in the vector. The VHH cDNAs are expressed as preys, with a N-terminal Gal4-activation domain fusion (Gal4ADZ70). A library of 2.1 million clones was obtained, collected and aliquoted. Tau variant 0N4R isoform (NM_016834.4) was expressed as bait with a C-terminal fusion with lexA (Tau-LexA) from pB29 vector, which is derived from the original pBTM116 (45). The library was screened at saturation, with 20 million tested diploids, using cell-to-cell mating protocol (46) by increasing the selection pressure with 100 mM 3amino-1,2,4-Triazol (3AT). The improvement in affinity of some selected clones was confirmed by one-to-one mating assay with L40ΔGal4 (mata) yeast strain transformed with the bait, corresponding to Tau variant 0N4R isoform with a C-terminal fusion with LexA (Tau-LexA) in pB29 plasmid, and Y187 (matα) yeast strain transformed with the prey, corresponding to a C-terminal Gal4-activation domain fusion (Z70-Gal4AD) (46). Diploids were grown on selection medium with increasing selection pressure by 3AT from 0 to 5 mM.

#### Production and purification of VHHs

Competent *Escherichia coli* BL21 (DE3) bacterial cells were transformed with the various PHEN2-VHH DNA constructs (13). For surface plasmon resonance (SPR) immobilization on chips and for crystallography assays, the VHHs were produced using a pET22b plasmid featuring a recombinant sequence encoding a pelB leader sequence, a 6-His tag, a TEV protease cleavage site and the considered VHH with or without an additional C-terminal cysteine. Recombinant *E. coli* cells produced proteins targeted to the periplasm after induction by 1 mM IPTG (isopropylthiogalactoside). Production was pursued for 4 hours at 28°C before centrifugation to collect the cell pellet.

Pellet was suspended in 200 mM Tris-HCl, 500 mM sucrose, 0.5 mM EDTA, pH 8 and incubated 30 min on ice. 50 mM Tris-HCl, 125 mM sucrose, 0.125 mM EDTA, pH 8 and complete protease inhibitor (Roche) were then added to the cells suspension and incubation continued 30 min on ice. After centrifugation, the supernatant corresponding to the periplasmic extract was recovered. The VHHs were purified by immobilized-metal affinity chromatography - IMAC (HisTrap HP, 1mL, Cytiva) followed by size exclusion chromatography (Hiload 16/60, Superdex 75, prep grade, Cytiva) in NMR buffer (50 mM sodium phosphate buffer (NaPi) pH 6.7, 30 mM NaCl, 2.5 mM EDTA, 1 mM DTT). For crystallography or SPR immobilization, VHHs were dialyzed against 50 mM Tris pH 8.0, 50 mM NaCl and cleaved with His-tagged TEV protease after a first IMAC step. TEV protease and cleaved 6-His tag were removed by a second IMAC step, the VHHs being recovered in the flow-through and concentrated to 400 to 500 µM for crystallography. Yield varies between the mutant VHHs, from around 1 mg (mutants 12, 15) up to 10 mg (mutant 20) per liter of bacterial fermentation.

#### VHH labeling and Western blotting

For Western blot experiments, 50 µM VHH with a terminal cysteine in NMR Buffer were first buffer exchanged against PBS supplemented with 0.1 mM TCEP using a Zeba Spin Desalting Column. VHHs were labeled using 250 µM 5(6)-TAMRA C6 maleimide (Euromedex) for 4 hours at 4°C. The reaction was quenched using 1 mM DTT and probe leftover were removed by buffer exchange against PBS using a Zeba Spin Desalting Column. Following SDS-PAGE, proteins were transferred from the polyacrylamide gel on a nitrocellulose membrane and blocked in PBS with 0.1% Tween 20 and 5% milk for 1 hour. Labeled VHHs were used at a final concentration of 100nM in PBS with 0.1% Tween 20 for 1 hour incubation followed by three washes in PBS with 0.1% Tween 20. Image acquisition was done using Amersham ImageQuant™ 800 (Cytiva).

#### Production and purification of labelled ^15^N Tau 2N4R

pET15b-Tau recombinant T7lac expression plasmid was transformed into competent *E. coli* BL21 (DE3) bacterial cells. A small-scale culture was grown in LB medium at 37 °C and was added at 1:10 V/V to 1L of a modified M9 medium containing MEM vitamin mix 1X (Sigma-Aldrich), 4 g of glucose, 1 g of ^15^N-NH_4_Cl (Sigma-Aldrich), 0.5 g of ^15^N-enriched Isogro (Sigma-Aldrich), 0.1 mM CaCl_2_ and 2 mM MgSO_4_. Recombinant ^15^N Tau production was induced with 0.5 mM IPTG when the culture reached an optical density at 600nm of 0.8. Proteins were first purified by heating the bacterial extract, obtained in 50 mM NaPi pH 6.5, 2.5 mM EDTA and supplemented with complete protease inhibitors cocktail (Sigma-Aldrich), 15 min at 75 °C. The resulting supernatant was next passed on a cation exchange chromatography column (Hitrap SP sepharose FF, 5mL, Cytiva) with 50 mM NaPi pH 6.5 and eluted with a NaCl gradient. Tau proteins were buffer-exchanged against 50 mM ammonium bicarbonate (Hiload 16/60 desalting column, Cytiva) for lyophilization. Detailed procedure can be found in (47).

#### Nuclear Magnetic Resonance Spectroscopy Experiments

Analysis of the ^15^N Tau/VHH interactions were performed at 298K on a Bruker Avance Neo 900MHz spectrometer equipped with cryogenic probe. TMSP (trimethyl silyl propionate) was used as internal reference. Lyophilized ^15^N Tau were diluted in NMR buffer with 10% D_2_O and mixed with a VHH at 100 μM final concentration for each protein. 200 μL of each mix in 3 mm tubes were sufficient to obtain the 2D ^1^H, ^15^N HSQC spectra with 32 scans. ^1^H, ^15^N HSQC were acquired with 3072 and 416 points in the direct and indirect dimensions, for 12.6 and 25 ppm spectral windows, in the ^1^H and ^15^N dimensions respectively. Data were processed with Bruker Topspin 3.6 and analyzed with Sparky (T. D. Goddard and D. G. Kneller, SPARKY 3, University of California, San Francisco). The intensity plots were designed with matplotlib (48).

#### Surface Plasmon Resonance experiments

Affinity measurements were performed on a BIAcore T200 optical biosensor instrument (Cytiva). Recombinant Tau proteins were biotinylated with 5 molar excess of NHS-biotin conjugates (Thermofisher) during 4 hours at 4 °C. Capture of biotinylated Tau was performed on a streptavidin SA sensorchip in HBS-EP+ buffer (Cytiva). One flow cell was used as a reference to evaluate unspecific binding and provide background correction. Biotinylated-Tau was injected onto a SA chip (Cytiva) at a flow-rate of 30 μL/min, until the total amount of captured Tau reached 500 resonance units (RUs). VHHs were injected sequentially with increasing concentrations ranging between 0.125 and 2 µM in a single cycle, with regeneration (3 successive washes of 1M NaCl) between each VHH. Single-Cycle Kinetics (SCK) analysis (49) was performed to determine association *k*_on_ and dissociation *k*_off_ rate constants by curve fitting of the sensorgrams using the 1:1 Langmuir model of interaction of the BIAevaluation software 2.0 (Cytiva). Dissociation equilibrium constants (Kd) were calculated as *k*_on/_*k*_off_.

#### Thermal Stability Assay

Assays were carried out using a concentration of 80 µM VHH in NMR Buffer and 10x SYPRO® Orange (Invitrogen). 20 µL of each solution were placed in PCR multi-well plates and subjected to a temperature ramp from 25°C to 94°C set at 1°C per 2 minutes in a MX3005P Real-Time PCR system (Agilent). Fluorescence is recorded every 2 minutes. Each test is carried out in triplicates. Thermal melting point (Tm) is defined as the maximal point of the derivative curve of the time evolution of the fluorescence intensity.

#### In-cell self-aggregation assay

HEK293 cells were seeded in 12-well plates (10^6^ cells per well); 24 h later, cells were transfected with plasmids encoding mCherry, or the different VHH-mCherry constructs together with lipofectamine in optiMEM, as recommended by the manufacturer (Invitrogen). Forty-eight hours later, medium was removed, and cells were washed in pre-warmed PBS before 30-min fixation at room temperature with 4% PFA. After three successive washes in pre-warmed PBS, the nuclei were stained with DAPI (1/10,000) for 15 min at room temperature. Cells were cover-slipped with VectaMount. Ten images per condition (n = 3 independent experiences) were acquired using a Zeiss AxioObserver Z1 (spinning disk Yokogawa CSU-X1, camera sCMOS Photometrics Prime 95B). The number of mCherry positive cells containing puncta was quantified from three independent experiments, 10 images per experiment and per group. Data were plotted using matplotlib and seaborn (50).

#### *In vitro* kinetic aggregation assays

Tau 2N4R aggregation assays were performed with 10 μM Tau and with increasing concentrations of VHHs (between 0 and 10 μM) in MES aggregation buffer containing 50 mM MES pH 6.9, 30 mM NaCl, 2.5 mM EDTA, 0.3 mM freshly prepared DTT, supplemented with 2.5 mM heparin H3 (Sigma-Aldrich) and 50 μM Thioflavin T (Sigma-Aldrich), at 37°C. Experiments were reproduced 3 times in triplicates for each condition. The resulting fluorescence of Thioflavin T was recorded every 5 min/cycle within 200 cycles using PHERAstar microplate-reader (BMG labtech). The measures were normalized in fluorescence percentage, 100% being defined as the maximum value reached in the positive Tau control, in each experiment. The bar plots are rendered with matplotlib.

#### Seeding assays in HEK293 reporter cell-line

Stable HEK293 Tau RD P301S FRET Biosensor cells (ATCC CRL-3275) were plated at a density of 100k cells/well in 24-well plates. At 60% confluency, cells were first transiently transfected with the various pmCherry-N1 plasmid constructs allowing expression of the mCherry-fused VHHs. Transfection complexes were obtained by mixing 500 ng of plasmid diluted in 40 µl of opti-MEM medium, which included 18.5 µL (46.25% v/v) of opti-MEM medium with 1.5 µL (3.75% v/v) Lipofectamine 2000 (Invitrogen). Resulting liposomes were incubated at room temperature for 20 min before addition to the cells. Cells were incubated for 24 hours with the liposomes and 1 ml/well of high glucose DMEM medium (ATCC) with Fetal Bovine Serum 1% (Life technologies). 8 µM of recombinant MTBD seeds were prepared *in vitro*, in the presence of 8 µM heparin, as described (35). Cells were then treated with MTBD seeds (10 nM/well) in the presence of transfection reagents forming liposomes as here above described.

#### FRET Flow Cytometry

Cells were recovered with trypsin 0.05% and fixed in 2% PFA for 10 min, then suspended in PBS. Flow cytometry was performed on an ARIA SORP BD (acquisition software FACS DIVA V7.0 BD, Biosciences). To measure CFP emission fluorescence and FRET, cells were excited with a 405 nm laser. The fluorescence was captured with either a 466/40 or a 529/30 nm filter, respectively. To measure YFP fluorescence, a 488 nm laser was used for excitation and emission fluorescence was captured with a 529/30 nm filter. mCherry fluorophore was excited with a 561 nm laser and fluorescence was captured with a 610/20 nm filter. To selectively detect and quantify FRET, gating was used as described (35, 51). The FRET data were quantified using the KALUZA software analyze v2 and an in-house python script using FlowKit (52). Three independent experiments were done at least in triplicate, with at least 10,000 cells per analyzed replicate. Data were plotted using matplotlib and statistics computed with SciPy (53).

#### VHH Z70 mutants crystallization and structure determination

Z70 mutants 1, 3 and 20 concentrated to 400-500 µM were incubated with 1 mM of PHF6 peptide (PGGGSVQIVYKPKK) for 30 minutes prior crystallization screening. From an initial screening of around 600 conditions, optimal crystallization conditions were found to be 1.7 M Ammonium Sulfate, 4.25% (v/v) Isopropanol, 30% (v/v) Glycerol for mutant and 3 1, or 0.095M tri-Sodium citrate pH 5.6, 19% (v/v) Isopropanol, 19% (w/v) PEG 4000, 5% (v/v) Glycerol for mutant 20 (both conditions found in the Cryos Suite, Qiagen). Crystals were evaluated at SOLEIL synchrotron beamline PX1 and PX2A. All crystals belonged to space group P6_5_22 with cell parameters suggesting that the asymmetric unit contains one monomer (98% probability estimated from Matthews coefficient). The best diffraction was obtained at a resolution of 1.83 Å for mutant 1, 2.35 Å for mutant 3 and 1.54 Å for mutant 20. Structure was solved using molecular replacement (MOLREP (54)) with pdb 7qcq as template and refined to a R_work_ of 0.21, 0.2 and 0.17, and R_free_ of 0.24, 0.25 and 0.21 for mutant 1, 3 and 20 respectively, using REFMAC5 (55) and COOT (56). The structures were deposited in the Protein Data Bank (PDB) with access code 8opi, 8pii and 8op0 for mutant 1, 3 and 20 respectively.

## Data availability

Raw data are deposited in zenodo database (https://doi.org/10.5281/zenodo.8083246) or can be shared upon request. Three-dimensional crystallographic structure coordinates and electron densities are available for mutant 1 at https://www.rcsb.org/structure/8OPI, for mutant 3 at https://www.rcsb.org/structure/8OPII and mutant 20 at https://www.rcsb.org/structure/8OP0.

## Acknowledgements

We thank Dr Z. Lens for support on the T200 biacore measurements and P. Legrand for valuable support during data collection at beamline PX1 at the SOLEIL synchrotron facility (Paris, France). We acknowledge SOLEIL for the provision of synchrotron-radiation facilities. We also thank the BiCel platform from US 41 - UAR 2014 – PLBS for microscopy and FRET data acquisition. The NMR facilities were funded by the Nord Region Council, CNRS, Institut Pasteur de Lille, European Union, French Research Ministry, and Univ. Lille. Financial support from the IR INFRANALYTICS FR2054 CNRS for conducting the research is gratefully acknowledged. This study was supported by the LabEx (Laboratory of Excellence) DISTALZ (Development of Innovative Strategies for a Transdisciplinary approach to Alzheimer’s disease ANR-11-LABX-01), by I-site ULNE (project TUNABLE) and by the ANR (project ToNIC, ANR-18-CE44-0016). Our laboratories are also supported by LiCEND (Lille Center of Excellence in Neurodegenerative Disorders).

## This article contains supporting information

### Conflict of interest

J.-C.R. is CEO of Hybrigenic services

Part of the results is included in patent WO2020120644 NEW ANTI TAU SINGLE DOMAIN ANTIBODY Landrieu I, Buée L, Dupré E., Danis C, Rain J.-C., Arial A.

### Credit author statement

**Justine Mortelecque:** Validation, Investigation, Writing - Review & Editing. **Orgeta Zejneli:** Investigation, Writing - Review & Editing. **Séverine Bégard:** Validation, Investigation. **Marine Nguyen:** Investigation. **François-Xavier Cantrelle:** Investigation. **Xavier Hanoulle:** Methodology, Writing - Review & Editing, Supervision. **Jean-Christophe Rain:** Conceptualization, Methodology, Writing - Review & Editing. **Morvane Colin:** Methodology, Formal analysis, Visualization, Writing - Review & Editing, Supervision. **Luc Buée:** Conceptualization, Methodology, Writing - Review & Editing, Supervision, Project administration, Funding acquisition. **Isabelle Landrieu:** Conceptualization, Methodology, Writing - Original Draft, Supervision, Project administration, Funding acquisition. **Clément Danis:** Conceptualization, Methodology, Investigation, Writing - Review & Editing, Supervision. **Elian Dupré:** Conceptualization, Methodology, Validation, Formal analysis, Investigation, Data Curation, Writing - Original Draft, Visualization, Supervision.

## Notes

https://doi.org/10.5281/zenodo.8083246

https://www.rcsb.org/structure/8OPI

https://www.rcsb.org/structure/8OPII

https://www.rcsb.org/structure/8OP0

