## Supplementary material for "Strategy of selection and optimization of single domain antibodies targeting the PHF6 linear peptide within the Tau intrinsically disordered protein": Fig. S1

| | | |
|---|---|---|
| R47I | S23C A64V | P101S G115E |
| R47I | S23C G115E | F13Y R47K E48A |
| R47I | A25S R47K | S23I R47K E56D |
| R47K | A42V G115E | S23C G58V G115E |
| R47K | K45N G115E | S23C G58V G115E |
| T96S | R47K S59L | S23C G58V G115E |
| T96S | R47K Y97C | S23N K79E G115E |
| P101S | R47I P101L | T32I E56G G115E |
| G115E | A64V G115E | R47K Y113N Q116H |
| G115E | D65V G115E | R90G P101S G114E |
| G115E | D65N G115E | A8G Q15R M36L R47I |
| G115E | D76V G115E | S27A N77S V82A T96S |
| Q15H N80T | N77K G115E | G58E N80T Y83S G115E |
| G17E G115E | S78P G115E | |
| S23C R47I | K79E N80T | |

**Fig. S1** List of 43 mutants obtained from the initial yeast two-hybrid screen.
