## Supplementary material for "Strategy of selection and optimization of single domain antibodies targeting the PHF6 linear peptide within the Tau intrinsically disordered protein": Fig. S2

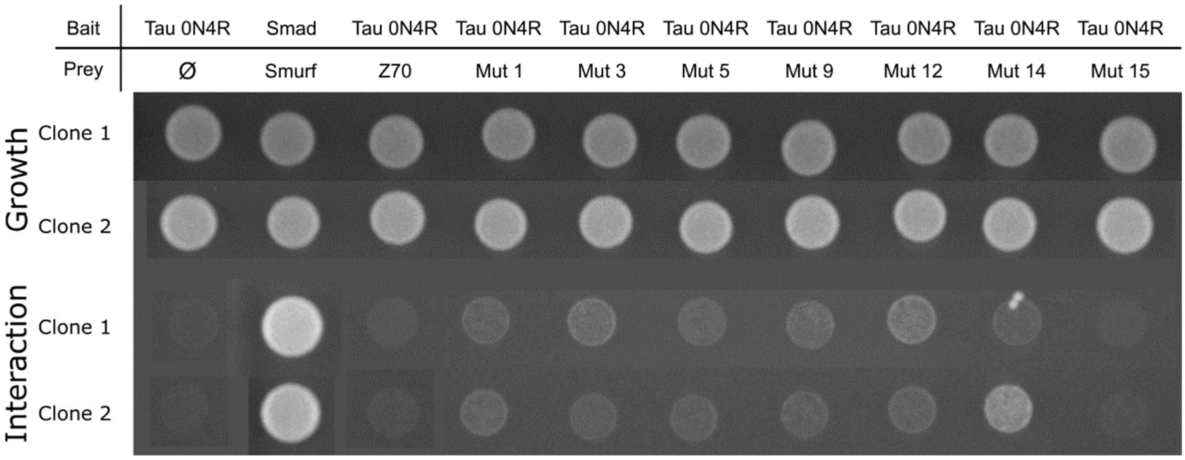


**Fig. S2** One-to-one mating by yeast two-hybrid on selective medium without tryptophane and leucine (growth) and without tryptophane, leucine and histidine and in the presence of 5 mM 3AT.
