## Supplementary material for "Strategy of selection and optimization of single domain antibodies targeting the PHF6 linear peptide within the Tau intrinsically disordered protein": Fig. S3

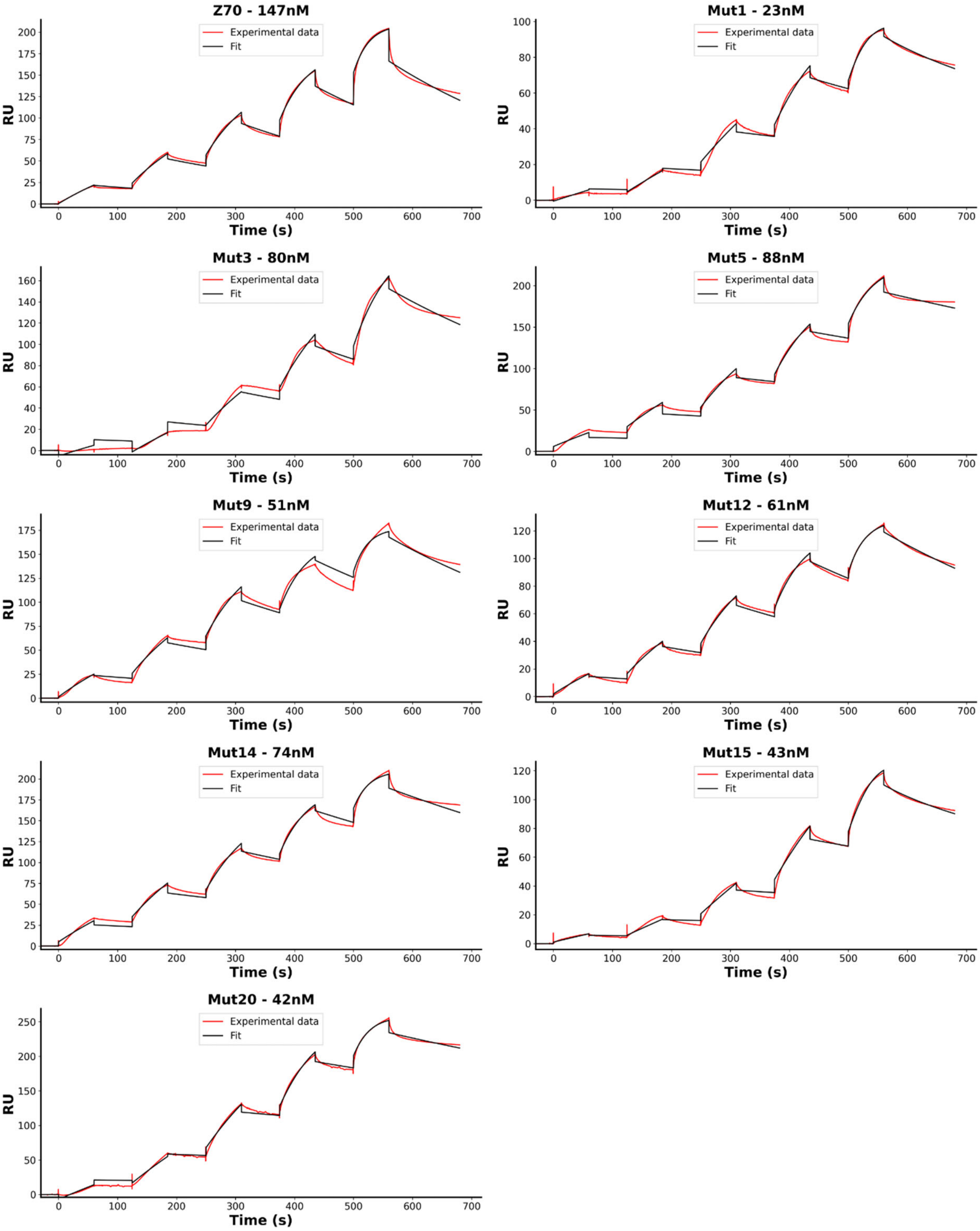


**Fig. S3** Sensorgrams of serial injections of VHHs (0.125, 0.25, 0.5, 1 and 2 μM) on a Tau immobilized SA chip in red and fitting with a 1:1 model in black.
