## Supplementary material for "Strategy of selection and optimization of single domain antibodies targeting the PHF6 linear peptide within the Tau intrinsically disordered protein": Fig. S4

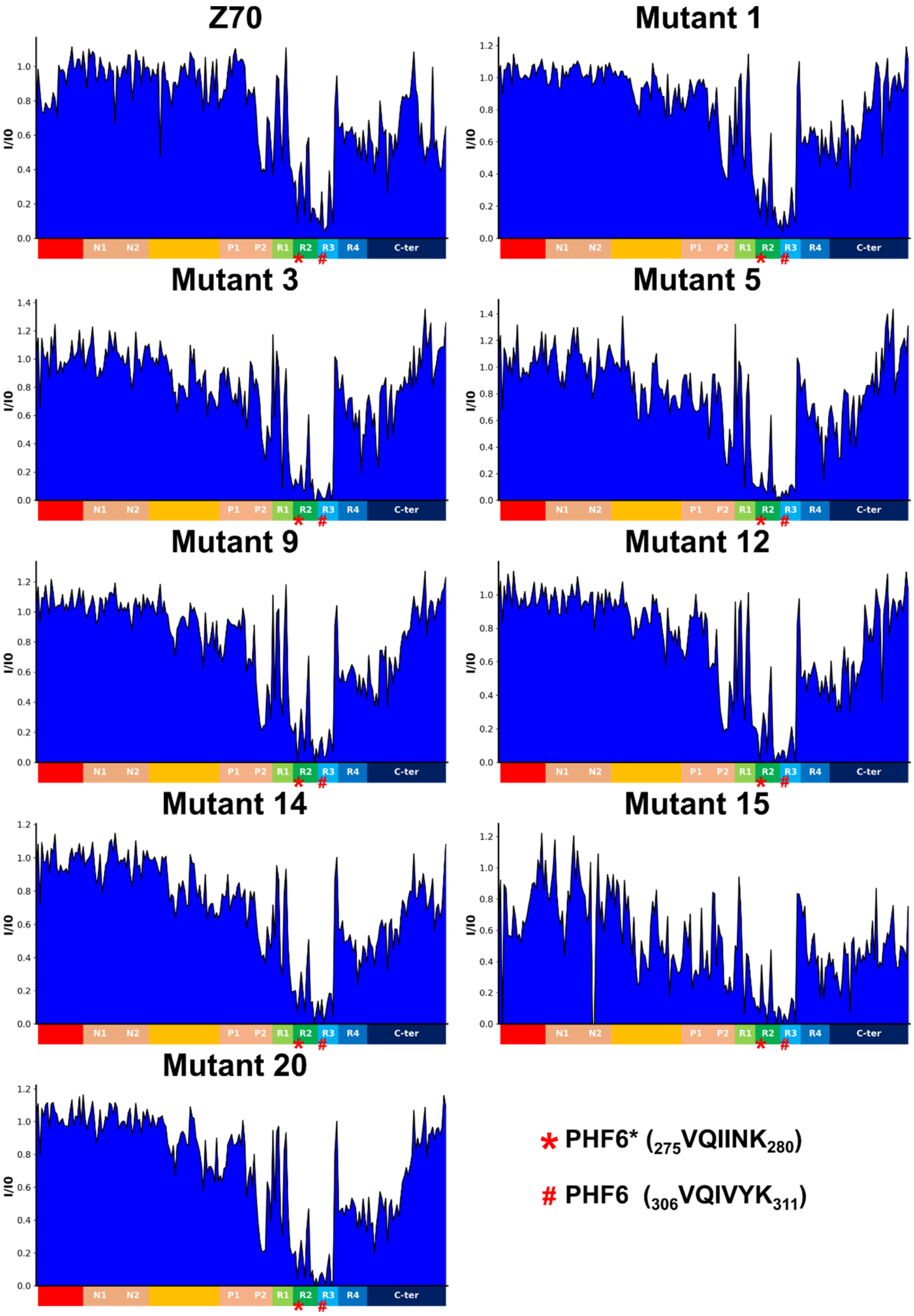


**Fig. S4** Normalized intensity ratios I/I0 of corresponding resonances in the two-dimensional spectra of Tau with equimolar quantity of the different VHHs (I) or free in solution (I0) for residues along the Tau sequence.
