## Supplementary material for "Strategy of selection and optimization of single domain antibodies targeting the PHF6 linear peptide within the Tau intrinsically disordered protein": Fig. S5

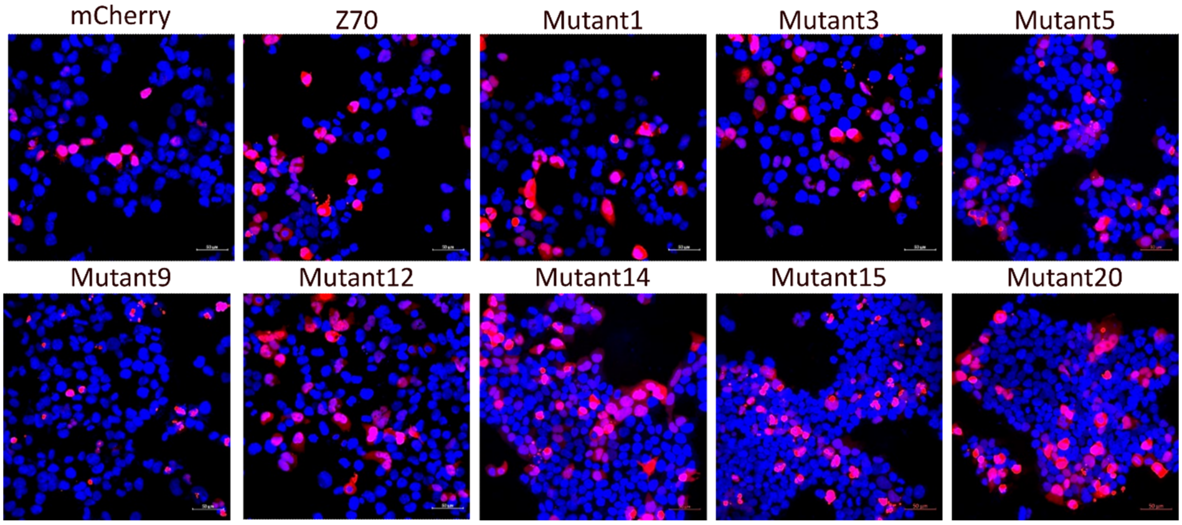


**Fig. S5** Self-aggregation of the mCherry constructs. The presence of aggregates is noticed by the appearance of "puncta" inside the cells in contrast with an uniform fluorescence across the cell in the absence of aggregates.
