## Supplementary material for "Strategy of selection and optimization of single domain antibodies targeting the PHF6 linear peptide within the Tau intrinsically disordered protein": Fig. S7

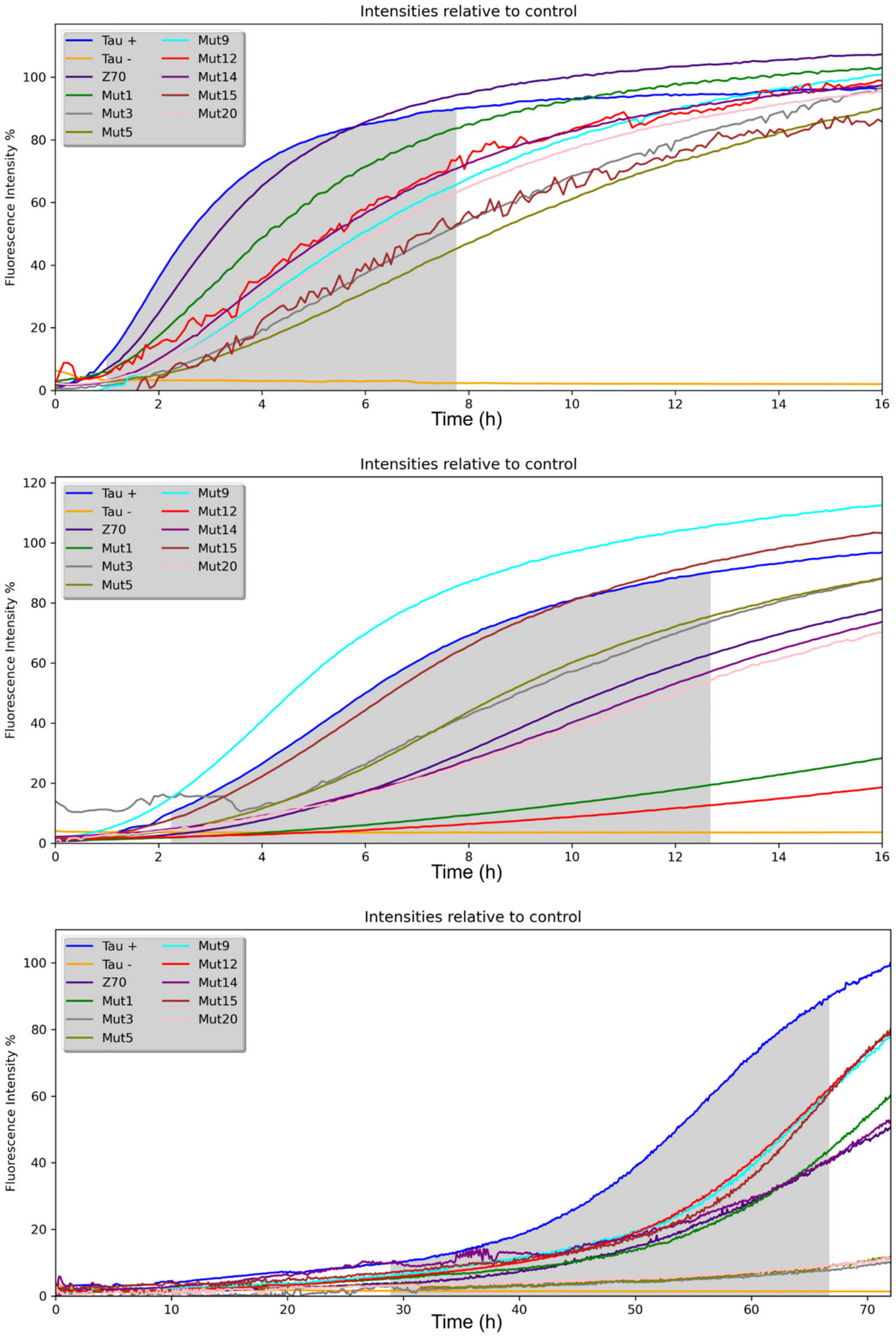


**Fig. S7** Intensity curves from the 0.2:1 VHH:Tau ratio *in vitro* aggregation assay from 3 independent experiments. The gray surface corresponds to 10-90% aggregation of the positive control which is used to compute inhibition efficiency of each VHH.
